# A stolen future – aberrant hippocampal neurogenesis produces glial cells in epilepsy

**DOI:** 10.1101/2024.09.23.614355

**Authors:** Segasby Toby, Sanaei Roozbeh, Aleksejenko Natalija, Mamad Omar, Henshall David, Floudas Achilleas, Heller Janosch

**Affiliations:** School of Biotechnology, Dublin City University, Dublin 9, Ireland; DCU Life Sciences Institute, Dublin City University, Dublin 9, Ireland; Department of Physiology and Medical Physics, RCSI University of Medicine and Health Sciences, Dublin 2, Ireland; FutureNeuro SFI Research Centre, RCSI University of Medicine and Health Sciences, Dublin 2, Ireland; Translational Immunology group, Medical School, University of Ioannina, Greece; Queen Square Institute of Neurology, University College London, Queen Square, London WC1N 3BG, United Kingdom; Biodesign Europe, Dublin City University, Dublin 9, Ireland

**Keywords:** Key-words: epilepsy, astrocyte, adult hippocampal neurogenesis, DCX, GFAP, single cell RNA sequencing

## Abstract

Adult hippocampal neurogenesis is disturbed in epilepsy. The increased neuronal activity in the epileptic brain leads to an increased production of newborn cells, increased mossy fibre sprouting and altered integration of new neurons within the hippocampus. Here, we set out to investigate increased astrocyte numbers following status epilepticus. We used immunolabelling of brain sections from the mouse intra-amygdala kainic acid model of epilepsy and publicly available single cell RNA sequencing datasets to assess newborn cells in the dentate gyrus. Similar to published series we found on increased number of reactive astrocytes present in the epileptic hippocampus. Additionally, we identified a cell population that expressed neurogenesis (doublecortin) and astrocyte (glial fibrillary acidic protein) markers in the epileptic brain, both in mouse and in human tissue. We further evaluated the expression profile of these cells. Immunolabelling showed expression of mature astrocyte markers aquaporin 4 and glutamate transporter-1. The single cell RNA sequencing data highlighted expression of neurogenesis and astrocyte markers in the doublecortin/glial fibrillary acidic protein-expressing cells. In conclusion, epilepsy pushes early neuroblasts to fate-switch to an astrocyte lineage as seen in the kainic acid-induced mouse model and in human resected brain tissue. Further understanding how neurogenesis is altered in epilepsy and whether the neuroblast fate-switch can be reverted will help in finding novel therapy strategies for epilepsy and other neurological diseases associated with aberrant adult hippocampal neurogenesis.

**Key Messages:** Following kainic acid-induced status epilepticus early neuroblasts appear to undergo a fate-switch to an astrocyte lineage. We were able to confirm the presence of cells positive for early neurogenesis and astrocyte markers in human epileptic tissue using scRNAseq data.

## Introduction

Adult hippocampal neurogenesis occurs within the dentate gyrus of the hippocampus. Here, neural stem cells generate immature neurons which ultimately differentiate into new granule cells^1^. This has been demonstrated in several animals but occurrence of adult neurogenesis in humans is still under debate^2–4^. Newly born immature neurons undergo a series of differentiation and migratory steps before integrating into existing circuits^1^. Adult hippocampal neurogenesis has attracted much attention due to its role in learning and memory^1,5^. Learning and exercise have been shown to increase the number of new neurons in the dentate gyrus^6–8^ and increasing neurogenesis rescues cognitive function in old age^5^.

In brain pathologies such as Alzheimer’s Disease newly generated neurons within the dentate gyrus can form aberrant connections further exacerbating the damage occurring in the brain^9^. Moreover, astrocyte changes and increased astrogliosis disrupt regulation of neural stem cells in the dentate gyrus^10^. Additionally, problems with adult hippocampal neurogenesis have been described in epilepsy^11–16^. Increased neuronal activity leads to aberrant production of newborn cells, increased mossy fibre sprouting and altered integration of new neurons within the hippocampus. These changes can drive the generation of seizures and establishment of epilepsy^17,18^.

Doublecortin (DCX) expression has been widely used as a marker for immature neurons. DCX is a microtubule-associated protein and is an essential protein facilitating migration of newborn neurons^19^. However, expression of DCX is not limited to newborn neurons. For example, its expression has been shown in astrocytes in the human neocortex^20^.

In addition to this, DCX-positive cells that resemble astrocytes morphologically and express glial fibrillary acidic protein (GFAP) have also been identified in epilepsy models^13,15,21^. A recent study has also shown that in physiological conditions, a small number of DCX-positive cells are not totally committed to neuronal differentiation and are able to regress and Text differentiate into astrocytes^21^. As seen in other studies, the type of chemoconvulsant used to induce epilepsy has an impact on how adult hippocampal neurogenesis is affected^11,13,15,21^. Although pilocarpine and kainic acid both increase the number of newborn cells, kainic acid specifically forces these cells into a glial cell fate^11,21^.

Here, we set out to investigate the DCX/GFAP-expressing cells in more detail as a first step to utilising manipulating adult hippocampal neurogenesis as a means to develop future epilepsy therapies. We confirmed the increased production of astrocytes in the intra-amygdala kainic acid model of epilepsy as well as the presence of DCX/GFAP-expressing cells in humans using publicly available single cell RNA sequencing databases.

## Subjects and Methods

### Experimental Model

All procedures were performed in accordance with the principles of the European Union Directive (2010/63/EU) and were reviewed and approved by the Research Ethics Committee of the Royal College of Surgeons Ireland (RCSI) (REC 842) under license from the Health Products Regulatory Authority (AE19127/I287). All efforts were made to minimise animal suffering and the number of animals used.

For the intra-amygdala kainic acid (IAKA) model of temporal lobe epilepsy (TLE), we used 10-week-old male C57BL/6OlaHsd mice originally from Harlan Laboratories (UK) and inbred at the Biomedical research Facility at RCSI. Induction of status epilepticus (SE) using the IAKA technique was performed as previously described^22^. Briefly, mice were anaesthetized (isoflurane; 5% induction, 1–2% maintenance) and equipped with a cannula (on the dura mater following coordinates from Bregma; IA: AP = −.95 mm, L = +2.85 mm, V = 3.1 mm).

Several days following implantation surgery, mice underwent IAKA microinjection (.3 μg KA in .2 μl; Sigma Aldrich Ireland) to induce status epilepticus, followed by an intraperitoneal (i.p.) injection of lorazepam (8 mg/kg) to ease convulsions and reduce mortality. Mice were allowed to recover in an incubator at 26°C.

Two weeks after kainic acid or PBS injection, mice were perfused transcardially with 4% paraformaldehyde (PFA), and brains were dissected out. Brains were placed in 4% PFA overnight at 4°C and then stored in PBS supplemented with 0.01% sodium azide at 4°C until sectioning.

All animals undergoing all procedures had ad libitum access to food and water.

### Immunohistochemistry

For sectioning, brains were embedded in 2% agarose and fixed to the vibratome submerged in 200mm glycerol. Sections were made at 30 mm intervals and stored in PBS supplemented with 0.01% sodium azide at 4°C until staining.

Sections were retrieved from both IAKA and PBS animals and placed into wells in staining pairs free floating. The tissue was first washed with PBS before quenching with 10 mM CuSO_4_ in 50 mM NH_4_Cl for 10 min. Sections then underwent successive wash steps first in ddH_2_O, then PBS before blocking in 3% BSA in 0.3% TBS+t (tris-buffered saline supplemented with 0.3% Triton X-100) for 3 hours at room temperature. The tissue was then incubated in 1% BSA in 0.1% TBS+t with primary antibodies DCX; 1:250 (ThermoFischer, CAT No. 481200, RRID: AB_2533840), GFAP; 1:1000 (ThermoFischer, CAT No. 14-9892-82, RRID: AB_10598206), AQP4; 1:250 (Synaptic Systems, CAT No. 429004, RRID: AB_2802156) and/or GLT1; 1:1000 (Merck, Cat No. AB1783, RRID: AB_90949) overnight at 4°C. The following day, tissue sections were washed in PBS and 0.1% PBS+t before incubating in secondary antibodies goat anti-guinea pig Alexa Fluor 488; 1:500 (ThermoFisher, CAT No A11073, RRID: AB_2534117), donkey anti-rabbit Alexa Fluor 594; 1:500 (ThermoFisher, CAT No. A21207, RRID: AB_141637), goat anti-mouse CF680; 1:500 (Sigma-Aldrich, CAT No. SAB4600361, RRID: AB_3492091) and Hoechst; 1:10,000 (Sigma-Aldrich, CAT No. 14533-100MG) in 1% BSA in 0.1% PBS+t at room temperature for 3 hours in the dark. Following further successive washing in 0.1% PBS+t, sections were mounted on glass slides with ProLong Diamond Antifade Mountant (ThermoFisher, CAT No. P36961).

### Image acquisition

Images were acquired on a Leica Stellaris laser scanning confocal with Power HyD S detectors, running Las X. Tile scans were performed with a 20x air objective for each hemisphere maintaining zoom factor, laser intensities and Z thickness with 37 1 μm step increments in 1024x1024 format. Tiles had 10% overlap and were statistically blended in the Las X software. Qualitative single frame images were acquired instead with a 100x oil objective with 37 0.75 μm step increments in 1024x1024 format.

All image analysis was performed using Fiji^23^. For fluorescence Intensity, ROIs were created for each layer of the dentate gyrus (Molecular Layer, Granule Cell Layer and Hilus, highlighted in Figure 1, A) using the Hoechst channel, with molecular layer boundaries confirmed with either AQP4 or GLT-1. ROIs were then applied to SUM slices Z-projections of each channel stack for mean grey value measurements. Cell counts were performed manually with Fiji cell counter for DCX expression and counter labels. Only cells with discernible cell body staining were counted.

**Figure 1:**
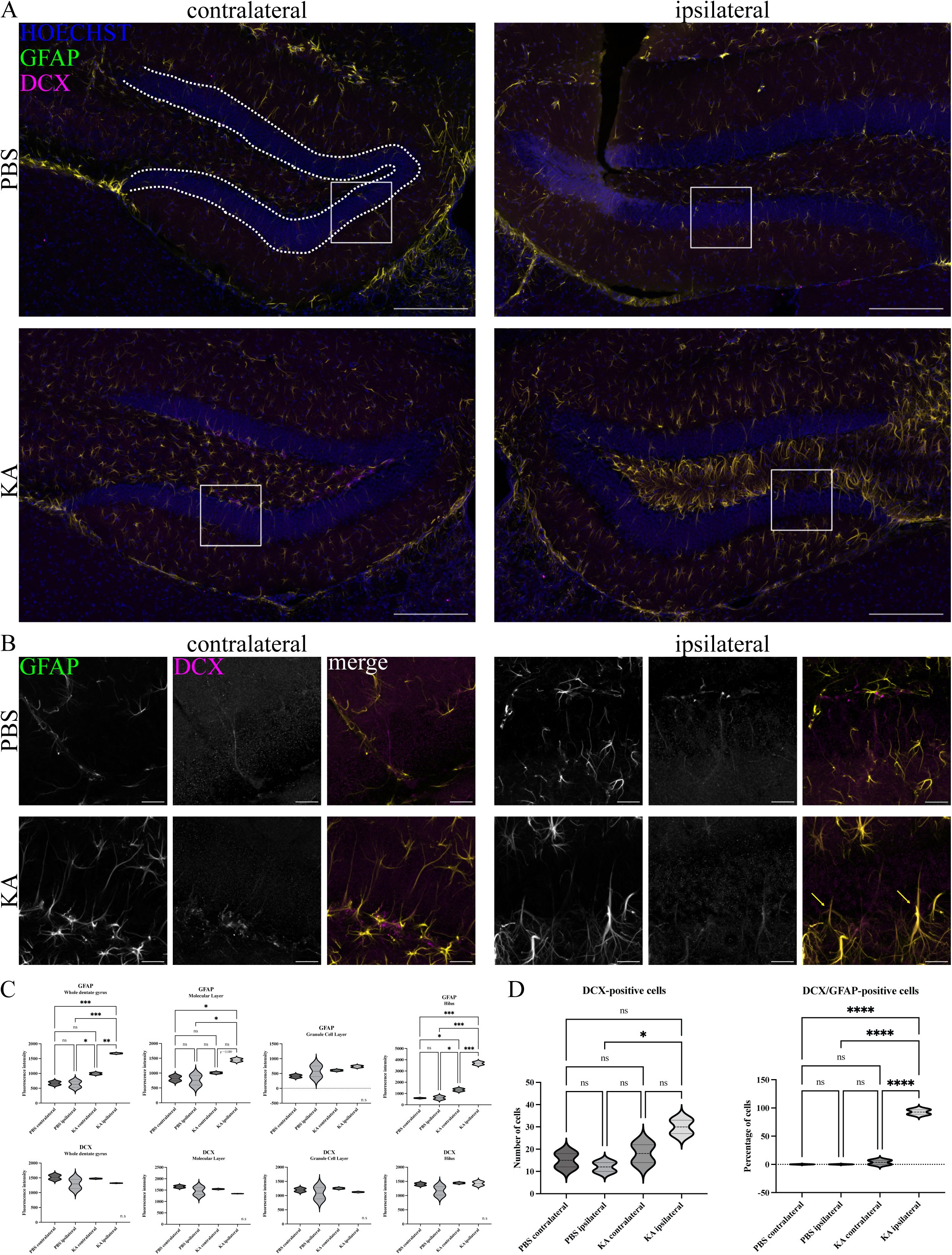
Expression of DCX and GFAP in the dentate gyrus in the IAKA model of epilepsy. A: Confocal images of PBS or KA-injected brains two weeks post SE. Sections were labelled with Hoechst (blue), GFAP (green) and DCX (magenta). Molecular layer, granular cell layer and hilus subregions are highlighted with white dashed lines. B: Area shown by white square in A at higher magnification. C: Fluorescence intensity measurements of GFAP (top row) and DCS (bottom row) in PBS or KA-injected brain sections. D: Cell counts of DCX-positive cells and cells positive for DCX and GFAP expressed as a percentage of DCX-positive cells. Significances are; *ANOVA*: * = *P* < 0.05, *** = *P* < 0.001 and **** = *P* < 0.0001. Scale bars = 200 µm (A) and 50 µm (B).

**Table 1:** Acquisition settings for quantification of confocal imaging. ^ All imaged, TauMode: Intensity, Operating mode: Analog, Gain 2.50---

| System |  |  | Single Frame Z stack |  | Tile Z Stack |  |
| --- | --- | --- | --- | --- | --- | --- |
| Format |  |  | 1024 x 1024 |  | 1024 x 1024 |  |
| Speed |  |  | 400 |  | 600 |  |
| Zoom factor |  |  | 1 |  | 2 |  |
| Image size |  |  | 116.82um x 116.82um |  | 58.12um x 58.12um |  |
| Pixel size |  |  | 114.19nm x 114.19nm |  | 56.82nm x 56.82nm |  |
| Optical section |  |  | 0.895um |  | 2.055 um |  |
| Pixel dwell time |  |  | 1.58us |  | 862ns |  |
| Frame rate |  |  | 0.063/s |  | 0.121/s |  |
| Line average |  |  | 1 |  | 1 |  |
| Frame average |  |  | 2 |  | 1 |  |
| Airy unit |  |  | 1 |  | 1 |  |
| Pinhole |  |  | 1 |  | 1 |  |
| Number of steps |  |  | 37 |  | 37 |  |
| Z-Step size |  |  | 0.750um |  | 1um |  |
| Tile overlap |  |  | - |  | 10% |  |
| Tile Blend |  |  | - |  | Statistical |  |
| Probe |  |  | 100x Objective |  | 20x Objective |  |
| Primary | Secondary | Excitation<br>wl | Laser<br>Power | Detector<br>Window | Laser Power | Detector<br>Window |
| Ms-GFAP | CF680 | 681 | 10 | 690-823 | 10 | 690-823 |
| Rb-DCX | AF594 | 590 | 30 | 597-650 | 30 | 597-650 |
| Gp-AQP4 | AF488 | 499 | 10 | 504-601 | 10 | 504-601 |
| Gp-GLT1 | AF488 | 499 | 15 | 504-601 | 15 | 504-601 |
| - | Hoechst | 405 | 2 | 425-500 | 2 | 425-500 |

### Statistical Analysis

All data comparisons between disease (KA) and control (PBS) groups as well as between layers of the DG were performed using One-Way ANOVA multiple comparisons tests. A p-value less than 0.05 was deemed statistically significant. The confidence interval was 95%. GraphPad Prism software (GraphPad Software Inc. by Domatics, CA, USA, version Prism 10.3.0) was used to perform all statistical analyses.

### Single cell RNA sequencing datasets

We performed secondary data analysis of single-cell RNA sequencing (scRNAseq) datasets - GSE140393^24^ and GSE209552^25^. We used the scRNA-seq data from five human epileptic patients with medically refractory temporal lobe epilepsy (GSE140393 dataset) and three temporal lobe tissue samples from non-neurological decedents as healthy controls (GSE209552 dataset).

### scRNA-Seq clustering and gene expression analysis

Using the Seurat package (v. 5.1.0) in R (v. 4.1), single cell reads from each sample were converted into Seurat objects, with data filtered so that genes detected in fewer than three cells for each object were excluded from downstream analyses. Cell doublets were identified and removed in each object using Doubletfinder (code available here: https://github.com/chris-mcginnis-ucsf/DoubletFinder). The SCTransform (v. 0.4.1) workflow was used to normalise and scale the data (code available here: https://satijalab.org/seurat/reference/sctransform); to normalise for differences in sequencing depth and stabilise gene-specific variances, sctransform removes unwanted variation by incorporating additional covariates and applies a log transformation to the data, resulting in an expression matrix suitable for downstream clustering and differential expression analysis. For clustering, individual objects were merged, normalised and scaled. Principle component analysis (PCA) was applied to the merged object to identify variable genes as input for clustering. Prior to clustering, a recently introduced function called IntegrateLayers (method=RPCAIntegration) was used to perform integration, which helps to co-embed common cell types and states across datasets using the cell’s PCA coordinates (code available here: https://satijalab.org/seurat/articles/seurat5_integration). Seurat’s FindNeighbors and FindClusters functions were used to identify the clusters, which were then visualised using a UMAP plot. Basic annotation was performed using key known markers of resident cells in the temporal lobe. Plotting the co-expression of GFAP and DCX markers of interest led us to target a cluster for further analysis. By subsetting this cluster, normalisation, scaling and PCA were performed followed by integration and clustering. We used the integration method harmony due to low cell numbers in cluster 11.

## Results

Aberrant adult hippocampal neurogenesis has been described in epilepsy^12,16,18^. The findings of most studies are based on Bromodeoxyuridine (BrdU) incorporation in dividing cells as well as the expression of DCX. Peculiarly, the expression of the immature neuronal marker DCX has been shown in other cell types, including astrocytes^20^. Aberrant proliferation and location of DCX-positive cells has been reported in rodent models of epilepsy, and also the presence of DCX- and GFAP-positive cells has been mentioned^13,15,21^.

First, we assessed DCX expression in tissue from the IAKA model (Figure 1). In agreement with previous studies, we saw an increase in GFAP expression highlighting astrogliosis and the presence of reactive astrocytes in the epileptic tissue (Figure 1). We found an increase in GFAP in the contralateral and the ipsilateral dentate gyrus in the IAKA model. To get a better understanding of the regions within the dentate gyrus that showed the highest upregulation of GFAP, we performed analysis of fluorescence intensities in the molecular layer, the granule cell layer and the hilus subregions (highlighted as dashed white line in Figure 1, A). The hilus displayed the largest increase in GFAP immunoreactivity, followed by the molecular layer (Figure 1, C). No significant difference in GFAP presence was found in the granule cell layer (Figure 1, C).

Additionally, we visualised the presence of DCX. When comparing fluorescence intensities across the different layers of the dentate gyrus, we did not see a difference between PBS and IAKA two weeks following SE (Figure 1, C). However, when counting DCX-positive cells there was an increase in the number of cells in our model in the ipsilateral dentate gyrus (Figure 1, D). Interestingly, when performing co-labelling of GFAP and DCX, we were able to identify a cell population of DCX/GFAP-expressing cells in the subgranular zone (Figure 1, D). We were not able to find any double positive cells in the PBS injected animals. However, the number of double positive cells was markedly increased in the IAKA model, with even a small number of DCX/GFAP-expressing cells present in the contralateral dentate gyrus (Figure 1, D).

To assess whether these cells are also present in human epilepsy, we took advantage of publicly available scRNA-Seq data. We used two data sets of human temporal lobe, one control [GSE209552]^25^ and one epilepsy [GSE140393]^24^ dataset. First, samples underwent pre-filtering followed by normalisation, scaling and integration to minimise the effects of technical sources of variation. Clustering resulted in 20 clusters in the control tissue (Figure 2A). None of the clusters were positive for DCX and GFAP together. Clustering of the epilepsy data set resulted in 16 distinct cell clusters (Figure 2, B). One of these clusters (cluster 11) showed co-expression of GFAP and DCX (Figure 2, B). The feature plots of the individual clusters revealed that cells of cluster 11 expressed glial cell markers EAAT2 (excitatory amino acid transporter 2, GLT-1, SLC1A2), OLIG2 (oligodendrocyte transcription factor 2) (Figure 2, C), neurogenesis markers GFAP, DCX, SOX2 (SRY (sex determining region Y)-box 2), ASCL1 (achaete-scute homolog 1), NCAM1 (neural cell adhesion molecule 1) (Figure 2, B and D), astrocyte markers S100beta, EAAT1 (GLAST, SLC1A3) (Figure 2, E), NG2 cell markers CSPG4 (chondroitin sulfate proteoglycan 4) and BEX1 (brain-expressed X-linked protein 1) (Figure 2, F), and neuronal markers MAP2 (microtubule-associated protein 2) and Thy1 (Figure 2, I). Cluster 11 was mostly devoid of cells expressing typical markers of microglial cells (Figure 2, G), endothelial cells (Figure 2, H) and immune cells (Figure 2, J). However, we also note the presence of a CD45-positive cell infiltrate consisting primarily of CD14-positive monocytes/macrophages and a small population of CD3-positive T-cells (Figure 2, J). Overall, cells in cluster 11 show expression features of reactive astrocytes.

**Figure 2:**
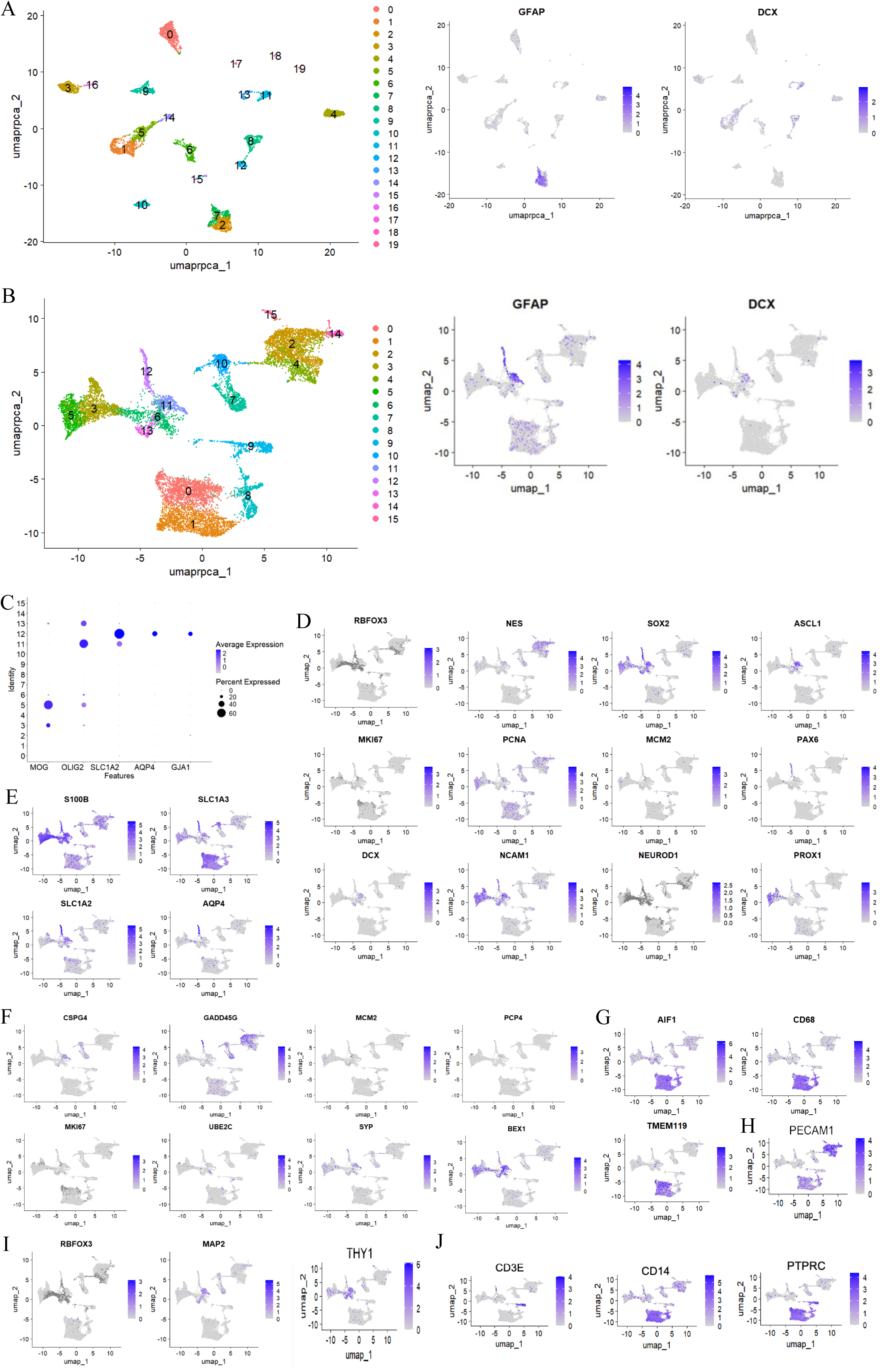
ScRNAseq data reveals increased number of DCX/GFAP-positive cells in human epileptic temporal lobe. A: UMAP plot of 20 clusters identified in the control temporal lobe tissue (left) and feature plots of GFAP and DCX expression (right). B: UMAP plot of 16 clusters identified in the epileptic temporal lobe tissue (left) and feature plots of GFAP and DCX expression (right). C: Plot highlighting glial cell marker expression across the 16 clusters identified in the epileptic temporal lobe tissue. D: Feature plots highlighting neurogenesis marker expression across the 16 clusters identified in the epileptic temporal lobe tissue. E: Feature plots highlighting astrocyte marker expression across the 16 clusters identified in the epileptic temporal lobe tissue. F: Feature plots highlighting NG2 cell marker expression across the 16 clusters identified in the epileptic temporal lobe tissue. G: Feature plots highlighting microglia marker expression across the 16 clusters identified in the epileptic temporal lobe tissue. H: Feature plot highlighting endothelial cell marker expression across the 16 clusters identified in the epileptic temporal lobe tissue. I: Feature plots highlighting neuronal marker expression across the 16 clusters identified in the epileptic temporal lobe tissue. J: Feature plots highlighting immune cell marker expression across the 16 clusters identified in the epileptic temporal lobe tissue.

Judging from the feature plots, cluster 12 possibly represents mature astrocytes not created through aberrant neurogenesis expressing important astrocyte proteins such as aquaporin 4, S100beta, EAAT1 (GLAST, SLC1A3) and EAAT2 (GLT-1, SLC1A2) and GFAP (Figure 2).

Lastly, we wanted to further explore whether the DCX/GFAP-positive cells were indeed functional astrocytes. In a first attempt we co-labelled brain sections with DCX, GFAP and essential markers of mature astrocytes, aquaporin 4 and EAAT2/GLT-1 (Figure 3, A). The DCX/GFAP-positive cells in the subgranular zone of the ipsilateral KA dentate gyrus did indeed express aquaporin 4 and EAAT2/GLT-1 (Figure 3, A).

**Figure 3:**
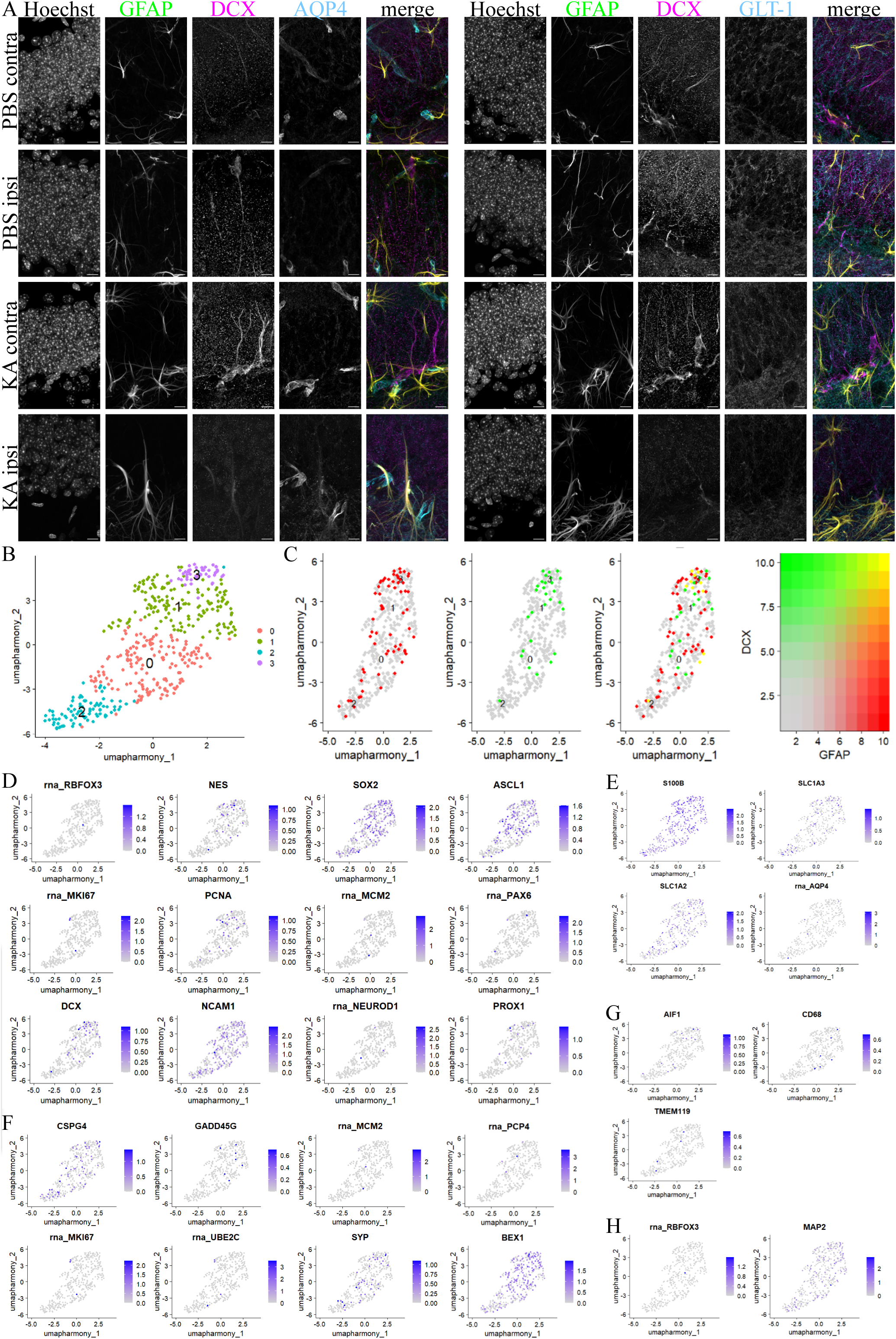
DCX/GFAP-positive cells express astrocyte markers. A: Confocal images of PBS or KA-injected brains two weeks post SE. Sections were labelled with Hoechst, GFAP (green), DCX (magenta) and aquaporin 4 (cyan, left) or EAAT2/GLT-1 (cyan, right). B: UMAP plot highlighting the four subclusters of cluster 11 in the epileptic temporal lobe tissue. C: Feature plot highlighting GFAP and DCX expression across the four subclusters of cluster 11 in the epileptic temporal lobe tissue. D: Feature plots highlighting neurogenesis marker expression across the 16 clusters identified in the epileptic temporal lobe tissue. E: Feature plots highlighting astrocyte marker expression across the 16 clusters identified in the epileptic temporal lobe tissue. F: Feature plots highlighting NG2 cell marker expression across the 16 clusters identified in the epileptic temporal lobe tissue. G: Feature plots highlighting microglia marker expression across the 16 clusters identified in the epileptic temporal lobe tissue. H: Feature plots highlighting neuronal marker expression across the 16 clusters identified in the epileptic temporal lobe tissue. Significances are; *ANOVA*: * = *P* < 0.05, *** = *P* < 0.001 and **** = *P* < 0.0001. Scale bars = 10 µm (A).

Additionally, we performed more in-depth expression analysis of the cells identified in cluster 11. The cells of cluster 11 were further subsetted (“isolated”), re-normalised, re-scaled and re-integrated to perform a higher resolution clustering to identify potential cellular subsets within cluster 11. Following this process, four distinct subsets of cells were identified within this cluster (Figure 3, B). Most DCX/GFAP-positive cells were found in subcluster 3 (Figure 3, C). As highlighted in the global expression plots in Figure 2, cells in cluster 11 expressed neurogenesis markers, astrocyte markers, NG2 cell markers, and neuronal markers but did not express typical markers of microglial cells (Figure 3, D-H). Subcluster 3 had the highest expression of DCX and GFAP with cells of cluster 3 co-expressing DCX and GFAP (Figure 2B).

## Discussion

Adult hippocampal neurogenesis is routinely altered under neuropathological conditions, and extensively demonstrated to be changed in TLE^12,16,18^. While the main output of adult hippocampal neurogenesis is neuronal, there are core glial contributions at all stages^1^. In particular astrocytes, the master regulators of the central nervous system are intimately involved in adult hippocampal neurogenesis and undergo extensive change across all neurological disorders including epilepsy^26,27^. Given the pathological shifts that occur in tandem between astrocytes and hippocampal neurogenesis this study sought to investigate their potential cross over.

Increased GFAP expression in epileptic tissue corroborates previous results^28^ exploring the effects of KA administration on the rate of gliosis. Astrogliogenesis is a common feature of epilepsy, characterised by astrocytic hypertrophy, functional dysregulation, disruption to regulatory gene expression and increased proliferation^27,29^.

There was expected to be a change in DCX expression in the KA model of TLE to concur an arrest of physiologically healthy adult hippocampal neurogenesis. Interestingly, DCX intensity measurements were comparable within each layer of the dentate gyrus across all experimental hemispheres in PBS and KA tissue slices. Of note however was the drastic change in DCX staining profile. Cells expressing DCX under confocal imaging in PBS and contralateral KA hemispheres are morphologically neuronal, with polarised processes from the lower granule cell layer out to the molecular layer. In KA ipsilateral dentate gyrus DCX signal is colocalised with GFAP and mature astrocyte markers aquaporin 4 and EAAT2/GLT-1. This staining profile was confirmed by cell counts which revealed co-expressing DCX and GFAP cells unique to epilepsy tissue. Previous work by Moura *et al.* eloquently showed this phenomenon^21^ and it is promising to see the result conserved with more readily available methodologies in a different model.

Reduced neuronal DCX is correlated with cell loss, reduced proliferation of hippocampal neural progenitor cells; or deviated differentiation in neuroblasts. Here, we demonstrate that 93% ± 6% of the elevated DCX cell counts in the ipsilateral dentate gyrus in the IAKA model were also GFAP positive. The reduction of neuronal output and favouring of astrogliosis is a distinct phenotype of TLE previously described^30^.

While DCX is indicative of differentiation stage and linage, the transition from progenitor cell to final cell fate can be modified^31^, with definitive determination not allocated until later stages. Astrocytes are a physiologically healthy differentiation end point in adult hippocampal neurogenesis^32^ all be it at much lower numbers than granule neurons. As astrogliosis is a fundament of TLE pathology^27,29^, and the neurogenic niche is sensitive to environmental changes, it could be possible that early neuroblasts are fate-switching to an astrocyte lineage in response to KA insult and frequent seizures as previously hypothesised^21^. This is consistent with the lack of a distinctive difference in DCX intensity measurements, but starkly changed cell morphologies and astrocyte marker co-labelling observed in epilepsy tissue.

However, GFAP is not an exclusive marker to astrocytes with radial glial neural stem cells in the dentate gyrus also expressing GFAP^33^. While fate-switching of neuroblasts is a corroborated pathway for explaining DCX/GFAP-positivity, alterative research demonstrates neural progenitor cells differentiation into astrocytes following quiescence in healthy aging^34^. This could represent another route by which TLE disturbs adult hippocampal neurogenesis processes and achieves greater GFAP levels throughout the sub granular zone.

It is crucial to be aware that GFAP expression patterns and neural stem cell proliferation, differentiation and survival are altered differently depending on chemoconvulsant administered^11,13,15,21^. The presence of GFAP and DCX-expressing cells in human single cell RNA-seq is particularly important here in defining the relevance of this potential stolen fate for human disease.

In conclusion, we demonstrated astrogliosis and a fate-switch of early neuroblasts to an astrocyte lineage in response to the KA insult. Additionally, we confirmed the presence of DCX/GFAP-positive cells in human epileptic tissue using scRNAseq data. These cells expressed glial cell markers. Future investigation is needed to fully characterise the DCX/GFAP-positive cells in human epilepsy and whether treating the aberrant production of glial cells can be used as therapy in epilepsy.

## Acknowledgement

This work was supported by funding from the Irish Research Council (EpiGlymph) under grant IRCLA/2022/3828 and by funding from Science Foundation Ireland and FutureNeuro industry partners (16/RC/3948, 21/RC/10294_P2). We thank Ms Oluwafadekemi Bello for performing pleminiary experiments.

